# Manipulation in grazing, viral pressure and resource availability leads to success in the isolation of abundant marine bacteria

**DOI:** 10.1101/2023.02.27.530246

**Authors:** Xavier Rey-Velasco, Ona Deulofeu, Isabel Sanz-Sáez, Clara Cardelús, Isabel Ferrera, Josep M. Gasol, Olga Sánchez

## Abstract

Isolation of microorganisms is a useful approach to gather knowledge about their genomic properties, physiology, and ecology, in addition to allowing characterization of novel taxa. We performed an extensive isolation effort on samples from seawater manipulation experiments that were carried out during the four astronomical seasons in a coastal site in the NW Mediterranean to evaluate the impact of grazing, viral mortality, resource competition and light on bacterioplankton growth. Isolates were retrieved using two growth media and their full 16S rRNA gene was sequenced to assess their identity and compute their culturability across seasons and experimental conditions. A total of 1643 isolates were obtained, which mainly affiliated to classes *Gammaproteobacteria* (44%), *Alphaproteobacteria* (26%) and *Bacteroidia* (17%). The most commonly isolated genera were *Alteromonas* and *Limimaricola*. While isolates varied across culture media, seasons and treatments, those pertaining to class *Gammaproteobacteria* were the most abundant in all experiments, while *Bacteroidia* was preferentially enriched in the treatments with reduced grazing. Sixty-one isolates had a similarity below 97% to cultured taxa and are thus putatively novel. Comparison of isolate sequences with 16S rRNA gene amplicon sequences from the same samples showed that the percentage of reads corresponding to isolates was 21.4% within the whole dataset, with dramatical increases in summer virus-reduced (71%) and diluted (47%) treatments. In fact, we were able to isolate the top-10 abundant taxa in several experiments and from the whole dataset.

**IMPORTANCE:** The traditional observation that we can only culture 1% of bacteria for a given environment has recently been questioned on several grounds, among other reasons because it is importantly influenced by environmental conditions. We cultured a high amount of heterotrophic bacterial strains from experiments where seawater environmental conditions had been manipulated and found that decreasing grazing and viral pressure as well as rising nutrient availability are key factors increasing the success in isolating marine bacteria. Our data clearly suggests that the “1% culturability paradigm” needs to be revised and reinforces bacterial cultures as a powerful way to discover new taxa.

## INTRODUCTION

Current marine microbial ecology is largely based on culture-independent studies, yet isolation of marine microbes is still an essential process that allows performing physiological experiments and testing ecological hypotheses derived from culture-independent studies, by allowing access to whole genomes that inform about microbial metabolic capabilities and characterization of novel genes (1), retrieval of novel taxa (2), and the utilization of naturally present organisms in *e.g.* bioremediation purposes.

The cultivation success of relevant marine microbes has been limited by the “Great Plate Count Anomaly” (3), the observation that only 0.001-1% of the cells or taxa from an environment seem to be culturable on agar plates. However, this paradigm has recently been challenged. Martiny (4) argued that 35% of ocean taxa had known cultured relatives with 97% similarity and that high proportions of bacteria are already culturable, although this figure has been lowered by Steen et al. (5) using data less skewed towards cultured organisms. Another study by Lloyd et al. (6) also reduced the percentages of taxa with cultured relatives. However, these studies collectively indicate that higher-than-thought proportions of taxa have been cultured across biomes, in particular in the oceans, questioning the “1% culturability paradigm”.

Various alternative culturing techniques have been developed in order to increase the retrieval in culture of microorganisms from environmental samples, such as diffusion chambers (7), cultivation chips (8), microfluidic systems (9), microencapsulation (10) or high-throughput dilution to extinction (11). Still, agar plates are the most economic and easy-to-implement method to culture microorganisms. This technique is biased towards copiotrophic taxa, fast-growing bacteria that generally are present in low abundances in the sea (12), among other reasons because they are especially targeted by protists and viruses (13–15). However, high culturabilities have been obtained using agar plates after sudden environmentally relevant events (16), or in nutrient-rich conditions (3, 17–21) suggesting that changes in environmental conditions could lead to a larger culturing efficiency. In fact, it is known that certain copiotrophs can quickly respond to changes in the environment, so that they sometimes dominate microbial communities (13, 22–24), and these environmental changes might be related to increased nutrient availability or reduced predator or viral mortality.

Micro- or mesocosm experiments are a common approach to determine the effects of environmental variables on microbial abundance, activity and diversity. They have been used *e.g.* to describe the effects of phytoplankton blooms (25), oil spills (26), grazer reduction (27) or viral suppression (14) on bacterial community dynamics using molecular approaches, but so far we are not aware of culturing efforts in this type of experiments. In this study, we performed extensive bacterial isolation efforts in several manipulation experiments carried out in the four astronomical seasons that evaluated the impact of grazers, viruses, light and resource availability on Blanes Bay Microbial Observatory (BBMO, NW Mediterranean) on bacterial community dynamics using two different culture media: the standard, nutrient-rich Marine Agar 2216 (MA) and Marine Reasoner’s 2A Agar (mR2A), which has lower concentration of nutrients. Our main objectives were: I) to obtain an heterotrophic bacterial collection from Blanes Bay as diverse as possible eventually retrieving novel taxa, II) to test the phylogenetic compositional differences of culturable bacteria across culture media, seasons and experiments, and III) to compare the isolates with 16S rRNA gene amplicon sequencing data from the same experiments to determine the influence of environmental conditions on culturability. Thus, we isolated bacteria from initial (t_0_) and final times (t_f_) of experiments where we manipulated seawater to remove large predators in light/dark cycles (predator-reduced light, PL) and in the dark (predator-reduced dark, PD), to increase nutrient availability through dilution of the original bacterial community while reducing predators in light/dark cycles (diluted light, DL) and to add to these manipulations virus reduction in light/dark cycles (virus-reduced light, VL). There were unmanipulated controls for these experiments in light/dark cycles (control light, CL) and in the dark (control dark, CD).

## RESULTS

### Composition and diversity of the isolate collection

We obtained 1643 bacterial isolates (listed in Table S1) belonging to 5 phyla, 7 classes, 24 orders, 52 families and 125 genera that clustered into 336 isolated operational taxonomic units (iOTUs, 99% clustering) and 715 zero-radius iOTUs (ziOTUs, 100% clustering). The number of isolates was relatively homogeneous across culture media (816 isolates in MA, 827 isolates in mR2A), seasons and treatments, and comparatively higher in t_0_ samples altogether (Table 1). The most abundant isolates pertained to classes Gammaproteobacteria, Alphaproteobacteria and Bacteroidia (Fig. 1A) and genera *Alteromonas* (360 isolates), *Limimaricola* (129 isolates), *Pseudoalteromonas* (111 isolates), *Bacillus* (78 isolates) and *Alcanivorax* (71 isolates). Interestingly, two of our isolates were affiliated to the recently described class Rhodothermia (28) and one to Verrucomicrobiae.

**Figure 1.**
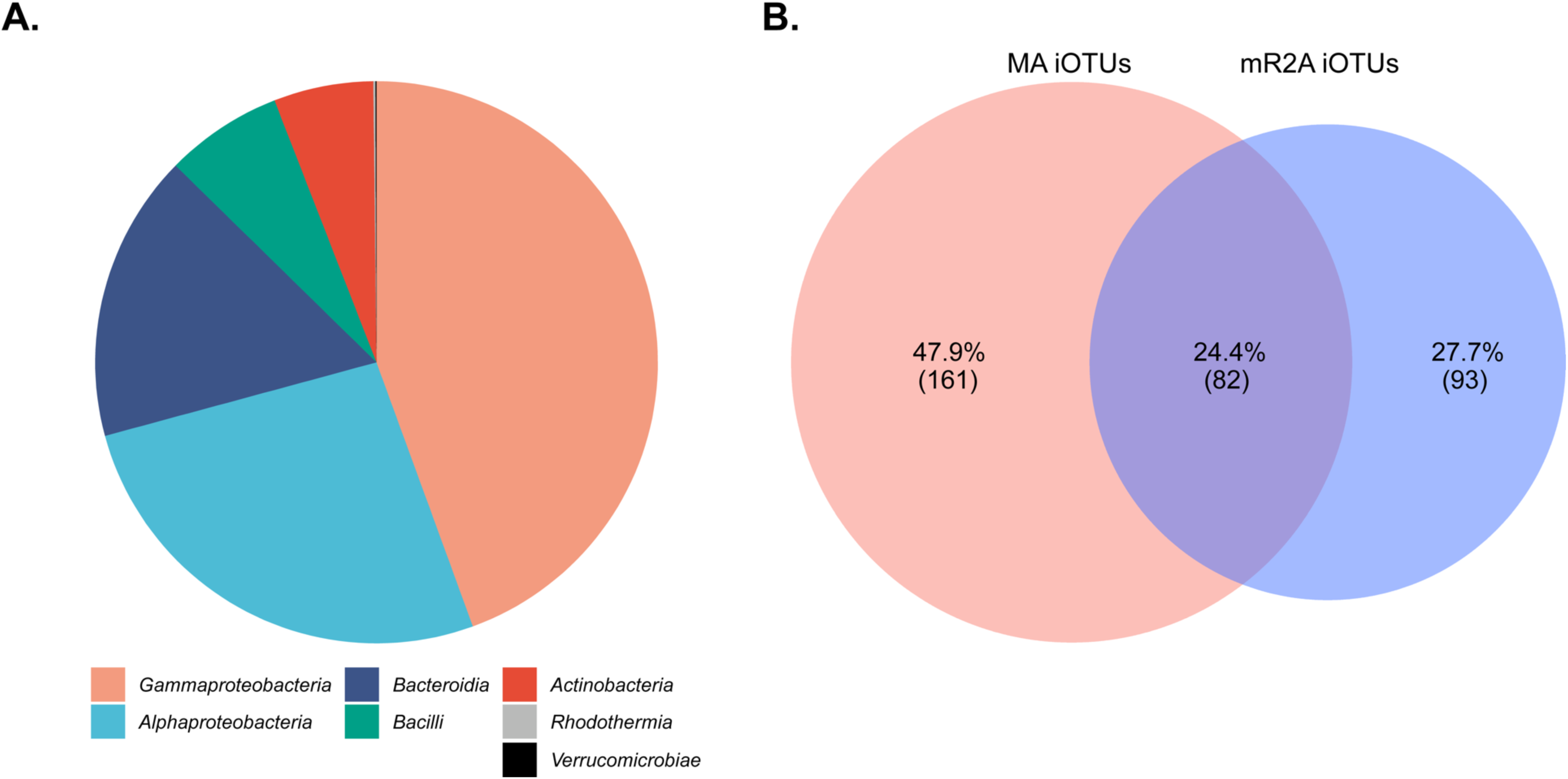
Overview of the isolate collection. **A.** Pie chart showing proportional class distribution of the iOTUs. **B.** Proportional Venn diagram showing similarity between iOTU composition in the two culture media used. Data calculated from the non-normalized iOTU table.

**Table 1.** Distribution of the isolates by class, season and treatment. . Relative abundances (in %) and number of isolates in brackets are presented for each class (based on SILVA classification).

| Class | Season |  |  |  | Treatment<sup>a</sup> |  |  |  |  |  |  |
| --- | --- | --- | --- | --- | --- | --- | --- | --- | --- | --- | --- |
|  | Fall | Winter | Spring | Summer | t<sub>0</sub> | CL | CD | PL | PD | DL | VL |
| *Gammaproteobacteria* | 42.6%<br>(156) | 45.9%<br>(206) | 29.5%<br>(117) | 58.1%<br>(251) | 33.8%<br>(194) | 39.6%<br>(61) | 35.9%<br>(56) | 55.5%<br>(96) | 52.0%<br>(78) | 56.2%<br>(122) | 56.2%<br>(123) |
| *Alphaproteobacteria* | 26.2%<br>(96) | 15.6%<br>(70) | 39.1%<br>(155) | 25.9%<br>(112) | 29.4%<br>(169) | 32.5%<br>(50) | 36.5%<br>(57) | 14.5%<br>(25) | 13.3%<br>(20) | 25.3%<br>(55) | 26.0%<br>(57) |
| *Bacteroidia* | 18.6%<br>(68) | 22.5%<br>(101) | 19.9%<br>(79) | 5.6%<br>(24) | 13.2%<br>(76) | 15.6%<br>(24) | 17.9%<br>(28) | 23.7%<br>(41) | 29.3%<br>(44) | 12.0%<br>(26) | 15.1%<br>(33) |
| *Bacilli* | 4.1%<br>(15) | 3.3%<br>(15) | 9.6%<br>(38) | 10.0%<br>(43) | 12.5%<br>(72) | 7.8%<br>(12) | 5.8%<br>(9) | 0.6%<br>(1) | 2.7%<br>(4) | 4.6%<br>(10) | 1.4%<br>(3) |
| *Actinobacteria* | 8.2%<br>(30) | 12.5%<br>(56) | 1.8%<br>(7) | 0.2%<br>(1) | 10.5%<br>(60) | 4.5%<br>(7) | 3.8%<br>(6) | 5.8%<br>(10) | 2.7%<br>(4) | 1.8%<br>(4) | 1.4%<br>(3) |
| *Rhodothermia* | 0.3%<br>(1) | 0.0%<br>(0) | 0.0%<br>(0) | 0.2%<br>(1) | 0.3%<br>(2) | 0.0%<br>(0) | 0.0%<br>(0) | 0.0%<br>(0) | 0.0%<br>(0) | 0.0%<br>(0) | 0.0%<br>(0) |
| *Verrucomicrobiae* | 0.0%<br>(0) | 0.2%<br>(1) | 0.0%<br>(0) | 0.0%<br>(0) | 0.2%<br>(1) | 0.0%<br>(0) | 0.0%<br>(0) | 0.0%<br>(0) | 0.0%<br>(0) | 0.0%<br>(0) | 0.0%<br>(0) |
| **Total** | 366 | 449 | 396 | 432 | 574 | 154 | 156 | 173 | 150 | 217 | 219 |

<sup>a</sup>t<sub>0</sub> considers all treatments at the initial time, CL to VL correspond to the final time of each treatment

The mean culturability measured as the ratio of the concentration of plate colony counts (CFU/mL) and total prokaryotic cell concentration (cells/mL) in DAPI (4′,6-diamidino-2-phenylindole) samples was 0.1 ± 0.22% and it almost always increased from t_0_ to t_f_, presenting its minimum in the fall (treatment PL at t_0_, mR2A, 0.001%) and its highest values in the fall VL t_f_ (the maximum was 1.4% in MA) and summer and spring DL and VL t_f_ (Table S2).

### (i) Isolates composition and diversity across culture media

In general, culturability (CFU DAPI ^−1^) had slightly higher values in MA, with a mean of 0.11 ± 0.24% than in mR2A, with a mean of 0.085 ± 0.2% (Wilcoxon Rank Sum test p = 0.07). The class composition of the isolates was highly similar in MA and mR2A, and Principal Coordinate Analysis (PCoA) did not show culture media to explain any compositional variation (Supplemental Figure S1A). However, while 51 genera were isolated with both media, 47 were only isolated in MA and 27 were unique for mR2A (Table S3). In fact, only 82 (24.4%) iOTUs were shared among MA and mR2A (Fig. 1B).

All α-diversity indices were significantly lower in mR2A than in MA (Figure S1B). Rarefaction curves of the different media showed a similar pattern: with an equivalent sampling effort we obtained more iOTUs on MA than on mR2A and the latter was closer to reach an asymptote (Fig. S1C).

### (ii) Isolate composition and diversity across seasons

The compositional comparison between seasons at class level is shown in Table 1. Gammaproteobacteria isolates had higher proportions in the summer experiment, while Bacteroidia were at their minimum. Alphaproteobacteria isolates were more numerous in spring, Actinobacteria were mostly isolated in winter and fall and the only isolate pertaining to class Verrucomicrobia was obtained in winter.

The isolates composition at the iOTU level was significantly affected by season as shown in the PCoA followed by *envfit* analysis (Pr[>r] < 0.001; Fig. S2A). A dendrogram of the iOTU table by season using Euclidean distances indicated clustering between winter and fall, with summer clearly separated (Fig. S2B).

Chao1 richness estimators and Shannon diversity indices of the isolated community were comparable in winter and fall, with lower values in spring and especially summer. Pielou evenness and Faith’s Phylogenetic Diversity (FPD) were lower in summer than in the rest of the seasons and had slightly higher values in the fall (Fig. S2C). Rarefaction curves indicated that with a similar sampling effort (slightly lower in fall) summer had almost reached the asymptote while winter, with the highest iOTU number, was far from it (Fig. S2D).

### (iii) Isolates composition and diversity across treatments

To compare treatments we clustered together all t_0_ samples (untreated) into one category and compared that category to each t_f_ (treatments). All treatments except controls were enriched in *Gammaproteobacteria* compared to t_0_. Alphaproteobacterial relative abundance increased slightly in the control treatments and decreased when predators were reduced. *Bacteroidia* were more abundant in predator-reduced treatments compared to the rest, while *Actinobacteria* and Bacilli were more abundant in t_0_ (*i.e.* reduced their presence in all experimental treatments). The rare classes, *Rhodothermia* and *Verrucomicrobiae*, were both isolated only in t_0_. On the other hand, light did not show any notable influence in isolates class composition (Table 1).

Regarding isolate similarity, the PCoA followed by *envfit* analysis showed that treatments explained the variance of iOTU composition with a significant goodness of fit (Pr[>r] < 0.001) with t_0_ samples opposed to PL, PD, DL and VL treatments, with t_f_ controls in the middle (Fig. S3A).

Due to the high variance of the t_0_ samples and the similarity of the light and dark treatments (Table 1), we only tested the α-diversity indices at t_f_ of light treatments. In general, isolates in DL and VL had the lowest values of the Chao1 estimator, Shannon diversity, Pielou evenness and FPD indices while CL and PL had similar, higher values (Fig. S3B). ANOVA and Tukey’s post-hoc test showed no significant differences between these values, probably due to the low number of samples in each category. Rarefaction curves (Fig. S3C, D) concur with this observation: among t_f_ DL and VL were the treatments with the least number of iOTUs despite being the most sampled.

To determine whether certain genera were associated to specific treatments, we carried out a Fisher’s Exact Test (Table S4). Among the 10 most abundant genera, *Bacillus*, *Alcanivorax*, *Erythrobacter* and *Palleronia* were significantly more likely to be isolated in t_0_ samples. *Alteromonas* was the most strongly associated genus to t_f_ of all treatments that implied manipulations and *Polaribacter* was related to all treatments except for CD. Apart from these genera, control treatments increased the possibility to isolate *Limimaricola*, *Tenacibaculum* and *Pseudoalteromonas*. Reduction of predators resulted in more probability to isolate genera *Pseudoalteromonas*, *Tenacibaculum* and *Dokdonia*; DL treatment was related with genera *Pseudoalteromonas*, *Limimaricola* and *Dokdonia*; and VL treatment was associated with *Limimaricola* and *Pseudoalteromonas*.

### Comparison between isolates diversity and amplicon 16S rRNA gene diversity

We compared the complete 16S rRNA gene sequences of our isolates with the V4-V5 region of the same gene of the amplicon sequence variants (ASVs) from the same experiments and we found that 63.08% of our ziOTUs matched to an ASV with 100% similarity. We isolated 173 out of 4594 ASVs (3.76%) accounting for 21.37% of the reads in the whole dataset, with high variability between samples: there were lower values in t_0_ samples with a mean of 2.44 ± 2.78% than at t_f_, with a mean of 9.97 ± 16.38% (Wilcoxon Rank Sum Test p < 0.01). Importantly, this value escalated in summer and spring VL and DL treatments (Fig. 4; Table S5 displays a complete list), reaching as high as 70.76% cultured reads in the summer VL treatment and 47.09% in spring VL. It also increased notably in the summer CD treatment, while in the fall and winter VL treatment the values were more modest but still higher than in t_0_.

While in most samples we cultured taxa pertaining to the rare biosphere (Fig. 5A, Fig. S4), in some ones we isolated very relevant ASVs in terms of abundance (Fig. 5B). We cultivated organisms that were 100% equal to almost all the top rank taxa detected by amplicon sequencing of the V4-5 region of 16S rRNA gene in summer VL and DL t_f_ treatments, affiliated to genera *Alteromonas*, *Vibrio* and *Limimaricola*. Interestingly, we also isolated ranks six and eight of summer CD t_f_ that pertained again to the genus *Alteromonas*. In the spring experiment, we cultured the first ASV of VL t_f_ and the second from DL t_f_, affiliating to genus *Nereida*. In the fall and winter experiments, we obtained more modest but still notable results from treatment VL t_f_. Ranks four, five and eight (genera *Alteromonas*, *Halomonas* and *Nereida*) were isolated in the fall experiment while in the winter experiment we cultured organisms 100% identical to ranks seven, eight, nine and ten (genera *Vibrio*, *Polaribacter*, *Lentibacter* and *Colwellia*). Surprisingly, there were some t_0_ samples from where some isolates were identical to relevant taxa: summer experiment, treatment DL t_0_ (rank three, *Limimaricola*) and fall experiment VL t_0_ (rank four, *Halomona*s). Importantly, we isolated organisms 100% identical to ranks three, six, seven and nine of the whole dataset pertaining to genera *Alteromonas* and *Vibrio*. Ranks, mean abundances and closest neighbors of all the ASVs with identical cultured organisms can be found in Table S6 for each season and treatment and Table S7 for the dataset as a whole.

### Novelty of the isolate collection

To test the novelty of our collection, we plotted the Closest Environmental Match (CEM) vs Closest Cultured Match (CCM) of our isolates (Fig. 2). This presents 61 isolates (3.7% of total isolates) that belong to 39 iOTUs (11.6% of total iOTUs) and 48 ziOTUs (6.7% of total ziOTUs) with less than 97% similarity with their CCM (*i.e.* they correspond to taxa that have never been cultured). Thirty isolates (1.8%) scored between 94.5 and 97 % similarity with both CCM and CEM, which could represent 19 genuinely novel species, as was revealed by clustering them to 97% similarity. Moreover, two isolates had less than 94.5% similarity with neither their CCM or CEM, and thus they could represent two novel genera (29).

**Figure 2.**
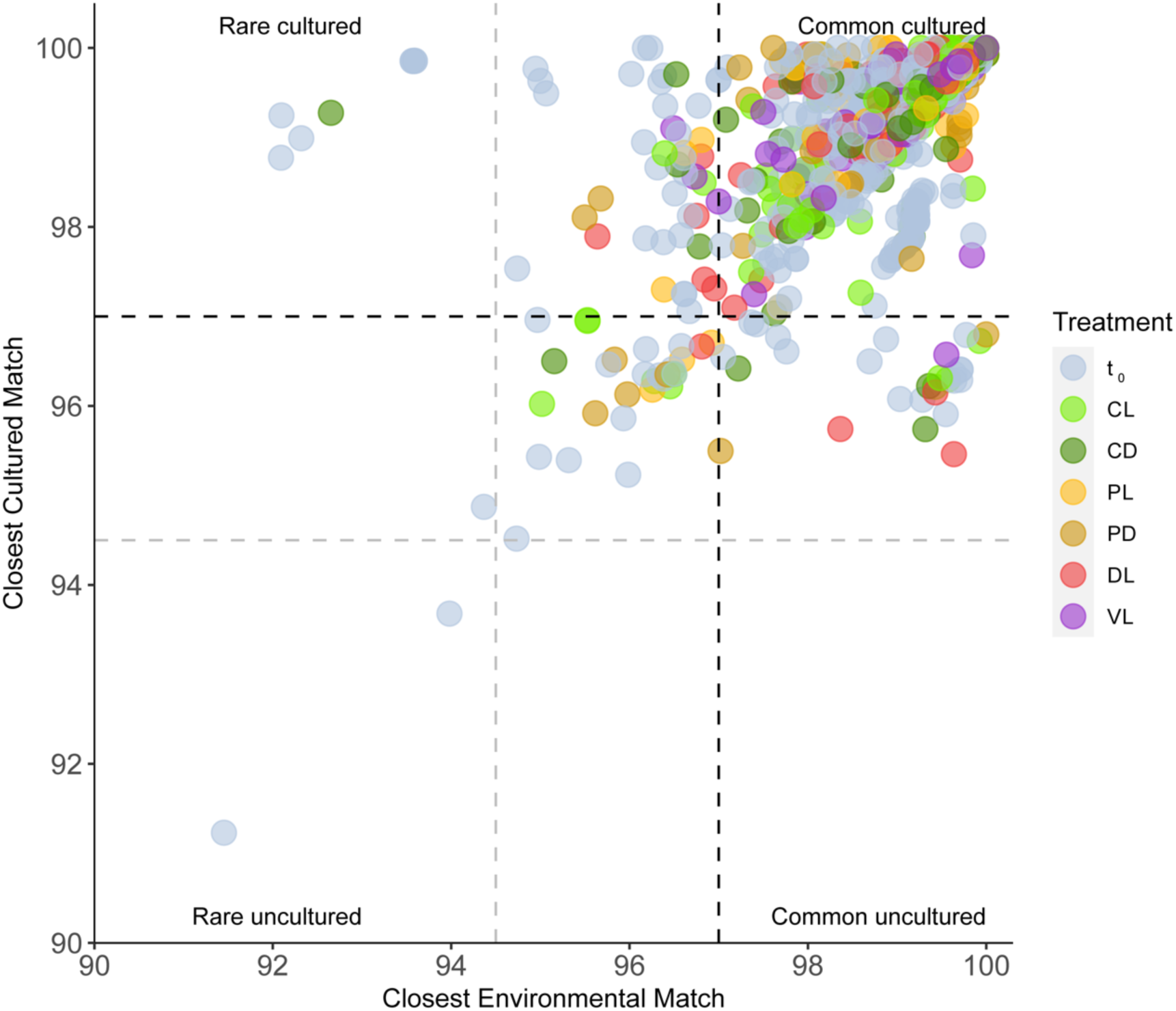
Percentage similarity between the CCM and the CEM of isolate 16S rRNA gene sequences. Horizontal and vertical lines represent the typical cut-off values of 97% (black dashed lines) commonly used for species delineation, and the cut-off values of 94.5% (grey dashed lines) used for genera delineation. Colored by treatment. Category t_0_ incorporates all treatments at the initial time; CL, CD, PL, PD, DL and VL correspond to the final times of each treatmen

To examine the phylogenetic placement of the novel strains, we constructed a phylogenetic tree with the putative novel ziOTUs (Fig. 3), which indicates that novel isolates pertained mostly to classes *Bacteroidia, Gammaproteobacteria and Alphaproteobacteria*. There were three ziOTUs affiliated to class *Bacilli*, and two ziOTUs to class *Rhodotermia*, which interestingly were the only ones from this class in the whole collection. Novel isolates were mostly cultured in MA (35 ziOTUs in MA, 13 in mR2A) and distributed similarly between seasons. Most novel ziOTUs were obtained from t_0_ samples (twenty-six) with some in control treatments (eleven), predator-reduced (nine), and less frequently DL (four) and VL (one) treatments.

**Figure 3.**
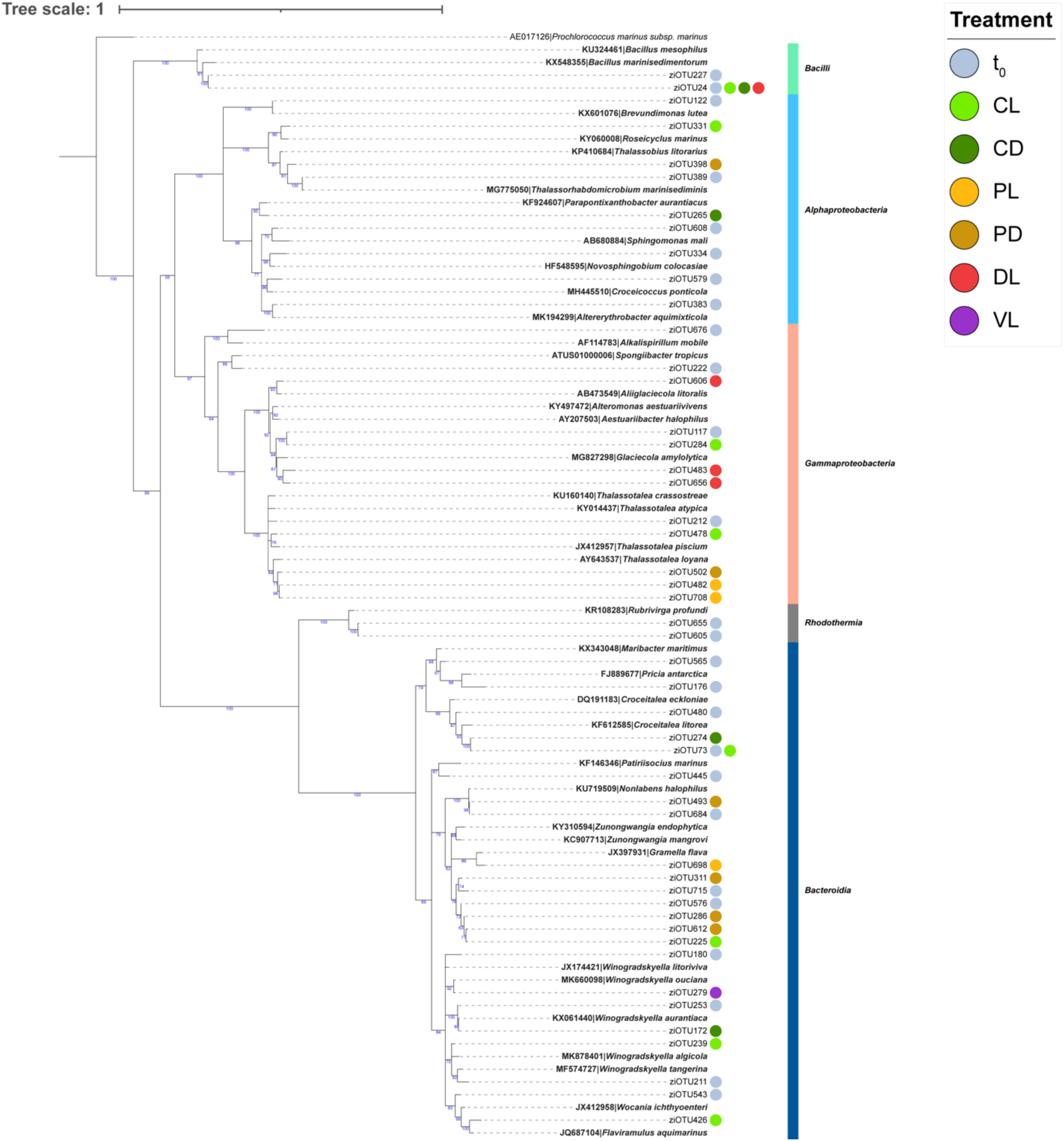
Phylogeny of putative novel isolates. Phylogenetic tree includes 48 ziOTUs with < 97% similarity to their CCM in the RDP database and their closest match in the SILVA Living Tree Project (in bold). The numbers in the nodes represent bootstrap coefficients calculated from 1450 replicates. Non-supported branches (bootstrap coefficients below 50%) were collapsed. Colored circles represent the treatments from where the ziOTUs were isolated. Classes to which the different taxa pertain are also indicated. Category t_0_ incorporates all treatments at the initial time; CL, CD, PL, PD, DL and VL correspond to the final times of each treatment.

## DISCUSSION

We have obtained an extensive collection of 1643 isolates from different manipulation experiments that were carried out in the four astronomical seasons, using two distinct culture media. Overall, the phylogenetic distribution of our collection at the class level (Table 1) is similar to that obtained in other studies made in the same sampling site (30). It is noteworthy, though, that classes *Rhodotermia* and *Verrucomicrobiae* had never been isolated in the BBMO, though the latter had been detected by culture-independent approaches (e.g. 30). In addition to this general picture, one of the aims of this study was to test differences in the phylogenetic composition and diversity of our isolates across culture media, seasons and treatments.

Culture medium MA has a higher organic matter concentration (5 g L^−1^ proteose peptone, 1 g L^−1^ yeast extract, total 6 g L^−1^ organic matter) than mR2A, while the latter presents a more diverse composition in terms of carbon sources (0.5 g L^−1^ proteose peptone, 0.5 g L^−1^ casamino acids, 0.5 g L^−1^ yeast extract, 0.5 g L^−1^ dextrose, total 2.5 g L^−1^organic matter), therefore the use of these two media aimed to increase the diversity of the obtained heterotrophic isolates. This goal was achieved, as the proportion of shared genera (Table S3) and iOTUs (Fig. 1B) across culture media was low. We expected to obtain higher CFU mL^−1^ in mR2A considering that previous studies have shown that more oligotrophic culture media resulted in better culturabilities than standard rich media (31, 32). However, in this study mR2A yielded slightly lower values than MA. This could have happened because mR2A is not low-nutrient enough to manifest this effect, and after all it still contains 2.5 g L^−1^ organic matter. Probably, using more nutrient-poor media such as the modified Seawater Medium (SW; 31, 33) could have led to the isolation of other taxonomic groups adapted to the low nutrients availability observed in the environment.

The isolate composition varied significantly across seasons: fall and winter were the most similar seasons and summer the most distant (Fig. S2B). This is the exact same trend that was observed for the structure of the environmental bacterial communities in the BBMO determined by DGGE Alonso-Sáez et al (34) and similar tendencies were observed by 16S rRNA gene amplicon sequencing (35). Our isolates reached their lowest α-diversity in summer with higher values in the spring and, especially, in the fall and winter (Fig. S2C). Other studies carried out in the BBMO based using molecular methodologies have shown similar tendencies (34, 35). While the seasonality of microbial communities in BBMO has been broadly described (34–38), this is, the first evidence of seasonality in the culturable bacteria in the BBMO.

The dominance of Gammaproteobacterial isolates in all treatments except the controls (Table 1) correlates with CARD-FISH relative abundances in these experiments (37) which were seen to be high for this class and especially *Alteromonadaceae* in the mentioned samples. Likewise, the isolation of comparatively more taxa affiliating to class *Bacteroidia* in the PL and PD treatments (Table 1) is in accordance with the same data: CARD-FISH abundance corresponding to this class peaked especially in the winter and spring PL and PD treatments (37). The PCoA by treatment (Fig. S3A) suggests that while the sole fact of confining seawater in bottles caused a change in the composition of culturable bacteria, a known effect that has already been reported (39, 40), treatments that implied a real manipulation of environmental conditions (PL, PD, DL and VL) had by themselves a notable effect in changing the composition of the culturable bacteria. This compositional change did not affect α-diversity in PL and PD, however in DL and VL treatments it decreased (Fig. S3B), suggesting that these treatments favored a narrow set of culturable bacteria over the other treatments. In the microcosm experiments performed by Teira et al (27), the α-diversity indices were reduced when reducing grazer pressure (equivalent to the PL treatment here), but here it did so only with deeper manipulations (DL and VL). While Teira et al. performed their experiments in offshore waters in different oceans (Atlantic, Pacific and Indian) that were analyzed with 16S rRNA gene amplicon sequencing, we observed a similar trend in the coastal Mediterranean sea.

When comparing the 16S rRNA gene sequences of isolates and ASVs (V4-V5 region), we observed that 63.08% of our ziOTUs were identical to an ASV and 3.76% of all ASVs were represented by isolates. It is common to find similar or lower proportions of isolates represented by sequences detected using molecular approaches (20, 30, 31, 41, 42). This is because taxa recovered by isolation usually belong to the “rare biosphere” of a given environment, while molecular techniques retrieve preferentially the relatively more abundant bacteria (12, 43). The low proportion of ASVs represented by isolates is actually similar to the value found in other studies (21, 41).

In this study we demonstrate that, using agar plates, it is possible to isolate dominant heterotrophic marine bacteria that represent a significant proportion of the population. In fact, our isolates accounted for 21.37% of the total reads and this percentage was higher in certain samples, reaching up to 70.76% in summer VL (Fig. 4; Table S5). Studies that compare isolates with sequencing data are scarce and they usually do not provide culturability values. Among the ones that have done this, Sanz-Sáez et al (20) obtained a mean of 0.3-1.1% with 7.8% as the maximum proportion of reads matching with isolates in samples from global oceans at different depths. Wang et al (21) isolated 45% of their whole 3-sample dataset from marine sediments, yet they applied a 97% similarity threshold to consider an OTU as cultured and thus, this number was likely an overestimation in comparison with our approach. Alejandre-Colomo et al (31) obtained a mean of 5.75 ± 2.99% cultured reads during a phytoplankton bloom in the North Sea, with a maximum of 11.51% in one sample (calculated from their Table S4), however they could not isolate any of the most abundant taxa. Sanz-Sáez et al (44) obtained isolates representing up to 45% of the reads in the deep ocean particle-associated fraction and isolated the most abundant ASVs in this fraction and ocean depth, reinforcing the idea that isolation of heterotrophic bacteria can be successful in certain environments or ecological situations.

**Figure 4.**
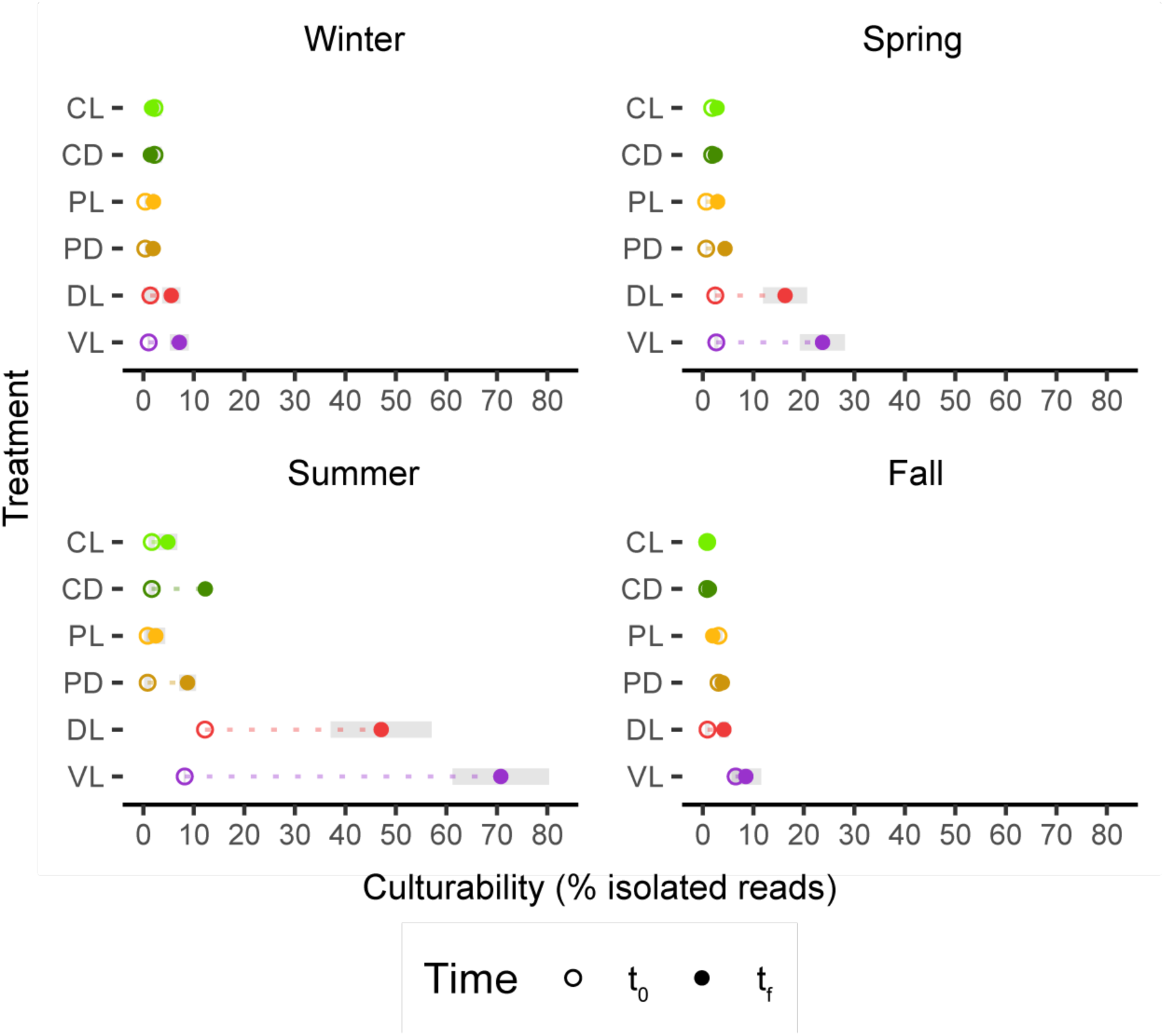
Cultured proportion of the population in each season, treatment and time. Percentage of Illumina 16S rRNA gene reads that corresponds to isolates at 100% identity vs treatments. Hollow circles represent t_0_, full circles represent t_f_. Grey areas indicate standard deviations of replicates.

**Figure 5.**
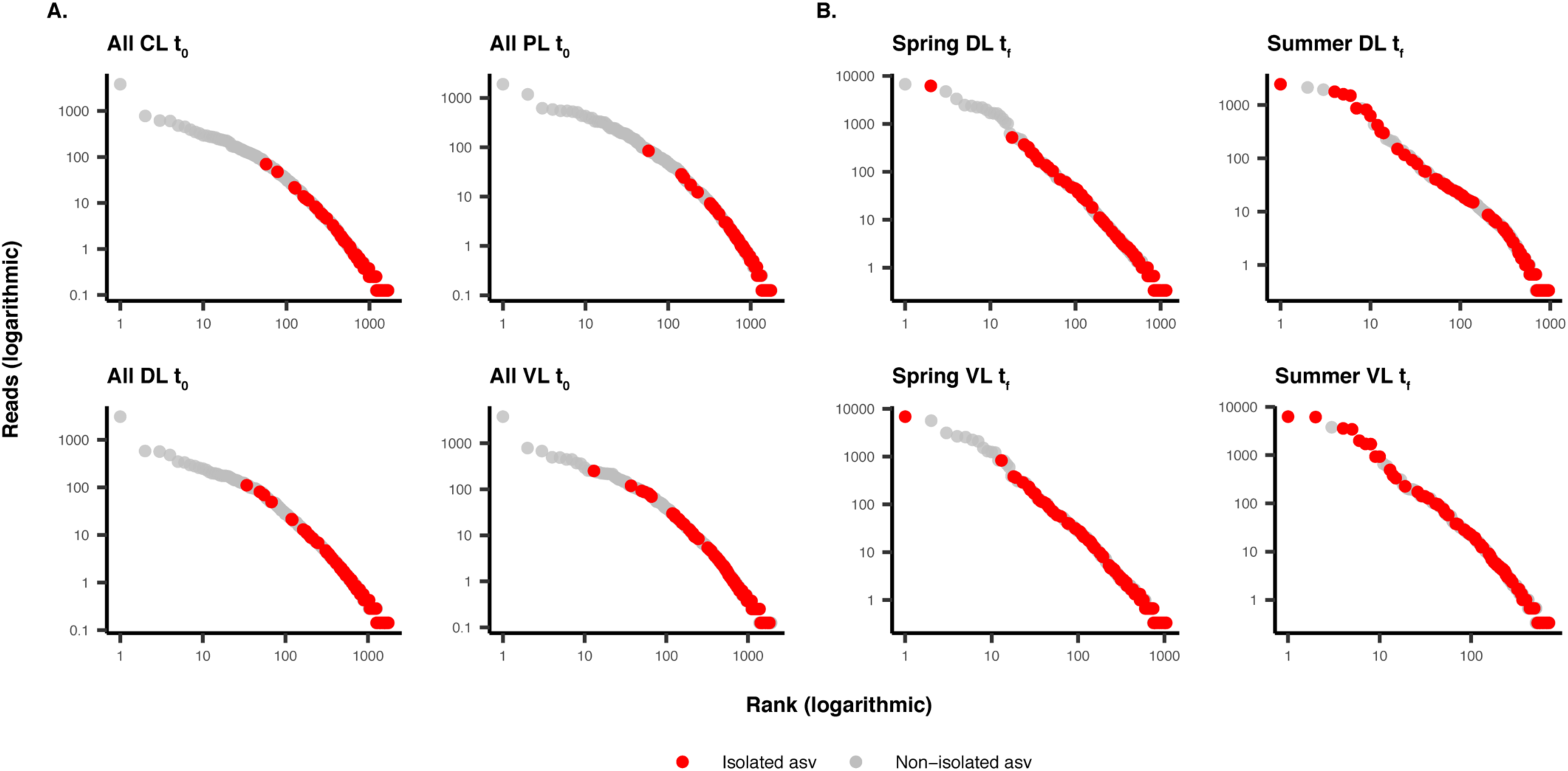
Selected rank abundance plots. based on 16S rRNA gene amplicon sequencing (region V4-V5) with indication of the cultured organisms that match the sequences (in red). The non-isolated ASVs are represented in grey. **A.** Examples where we only isolated rare taxa. All seasons are considered. **B.** Summer and spring samples where we isolated dominant taxa

In general, the DL and VL treatments were the ones with the highest proportions of reads in the 16S rRNA gene amplicon sequencing dataset matching exactly with an isolate (Fig. 4). To study this feature in more detail, we identified the ASVs that accounted for high percentages of reads in these treatments and that were 100% identical to isolates in this collection (Tables S6 and S7), mostly pertaining to genera *Alteromonas, Vibrio, Limimaricola* and *Nereida*. These genera are known fast-growing r-strategists that had commonly been isolated in other studies (20, 30, 31), and affiliate to phylogenetic groups which have been reported to actively respond to the increase of resource availability in the environment during events like phytoplankton blooms (23, 25, 45–47) or, in this case, treatments DL and VL (that provided increased resource availability). Copiotrophic taxa are known to be especially subject to grazer control and targeted by viruses (13–15), thus their growth and culturability is expected to be favored in the experimental manipulations in which these pressures were reduced. In fact, *Alteromonas* and *Limimaricola* were among the 10 most abundant isolates in this study and both genera appeared to be more frequently isolated in the DL and VL treatments according to the results of a Fisher’s Exact Test (Table S4). In view of these data, it looks like the high proportions of reads affiliating to cultured taxa in this study are caused by the effects of the experimental manipulations, especially DL and VL, that favored the growth of copiotrophic taxa which are normally found in the environment as part of the rare biosphere (12) but can become dominant under certain conditions, such as the ones created in the microcosms (14, 24, 48, 49).

It is noteworthy that the effect of experimental treatments in terms of isolation success appears to be markedly different between seasons. Summer was clearly the most successful season, followed by spring, while in the fall and winter experiments, the various treatments had very weak effects (Fig. 4; Fig. S4). This could be partially explained by looking at the identities of the most abundant ASVs that we did not isolate in this study with our methods of isolation. In the DL and VL treatments (t_f_), the most abundant uncultured taxa accounting for a high proportion of fall, winter and spring reads affiliated to a total of 12 species in the NCBI 16S rRNA database (Table S8) which had been isolated in different culture media and/or from other environments than the ones in this study (see references in Table S8). Only three of them had originally been isolated in MA from seawater: *Donghicola eburneus* (50), *Glaciecola amylolytica* (although it was isolated at 30°C; 51) and *Altibacter lentus* (isolated from 2000-m deep seawater; 52). It is also important to note that the cultures here were incubated at room temperature (RT; 20-25°C) and summer and the fall were the seasons with the closest *in situ* temperature to RT (23.1°C in summer, 19.5°C in fall; 37). All this information suggests that isolation success was higher in summer and spring because dominant taxa in these seasons were easier to isolate with our methods, especially our culture media. Moreover, the rarefaction curves by season (Fig. S2D) indicate that summer was precisely the best-sampled season followed by spring, so sampling effort could have also played a role.

It is well known that PCR-based sequencing overrepresents the abundances of bacteria with multiple 16S rRNA gene copies (53). In fact, all our isolates that accounted for high numbers of reads harbor numerous copies of the ribosomal operon according to rrnDB v. 5.8 (54), therefore it is likely that our proportions of reads corresponding to isolates are overestimated. For this reason, we generated a revised ASV abundance table by dividing each ASV read value by the number of 16S rRNA gene copies according to their class obtained from rrnDB v. 5.8 (54), and recalculated proportions of cultured reads according to these “corrected” values. With “correction”, the mean proportions of reads corresponding to isolates changed from 6.20 ± 12.23% to 5.48 ± 10.70% and the raw maximum of 70.76% decreased to 59.66% (Table S5). These estimations would only slightly reduce our isolation success, showing nonetheless the same trend.

The other main outline of this study is that our isolation effort has resulted in a high proportion (3.7% of all isolates) of putative novel taxa (Fig. 2). This novelty is comparatively higher than that found in other studies focused on isolation, such as the one from Ma et al (55), in which 1.9% isolates represented potential novel species in samples from deep sea water and sediments from the Mariana Trench using 6 culture media, or the 0.2% isolates representing novel genera in samples from different depths across the global ocean plankton using one culture medium (20). This could be explained as a result of the extensive isolation from a single sample from one site and also by the colony-picking strategy seeking different morphologies. It is noteworthy that the rarefaction curve of the site of study (Fig. S1D) had not reached an asymptote, implying that a greater isolation effort could have resulted in the discovery of more novel isolates. In fact, higher numbers of potential novel isolates have been obtained with a remarkable culturing effort using high-throughput isolation techniques (31). The novelty obtained in this study was not equally relevant across culture media as more novel ziOTUs were found in MA, which is logical if we attend to the alpha-diversity values (Fig. S1B) and rarefaction curves (Fig. S1C): MA was less selective than mR2A and permitted the culturability of a wider range of bacteria. It is also noteworthy that most putative novel taxa were isolated in t_0_ and control treatments. Taking into account that the sampling effort at t_0_ (considering all samples) was 3-4 times higher than that of the treatments at t_f_ (Table 1), it is not surprising to find more novel taxa there. Also, it is coherent that the frequency of isolation of novel taxa diminishes as the manipulation of environmental conditions is stronger, since it implies the selection of a smaller set of taxa. Alpha-diversity (Fig. S3B) and rarefaction curves (Fig. S3C, D) are in accordance with this idea: among final times, less diversity was found in more manipulated treatments (DL and VL).

We conclude that our extensive isolation effort applied to manipulation experiments resulted in the isolation of dominant taxa, corresponding to exceptionally high proportions of the microbial population, although they were mainly common copiotrophs. We are far from being able to obtain all the natural environment microorganisms in culture, but our data show that we can find a reasonable number of the bacteria responding to manipulations, indicating that culturability is highly influenced by environmental factors, especially resource availability, grazing and viral lysis.

Overall, our results further question the 1% culturability paradigm and point to environmental conditions as key factors influencing isolation success. Also, culturing microorganisms with traditional methods proves useful to discover novel taxa and isolate the most abundant members of the community under certain conditions.

## MATERIALS AND METHODS

### Origin of samples

Surface seawater samples were collected from the BBMO in the NW Mediterranean (41°40’N, 2°48’E), about 70 km north of Barcelona, and approximately 1 km offshore. Samples were collected on the four astronomical seasons: winter (21 February 2017), spring (26 April 2017), summer (5 July 2017) and fall (7 November 2017) and water was filtered *in situ* through a 200-μm mesh and transported to the laboratory within 2 h.

### Manipulation experiments

Six experimental treatments were set up the following day for each season as described in Sánchez et al (37). Briefly, the treatments consisted on: (i) unfiltered seawater in light/dark cycles (CL) and in the dark (CD), (ii) seawater prefiltered through a 1-μm filter to remove large predators while preserving most bacteria in light/dark cycles (PL) and in the dark (PD), (iii) unfiltered seawater diluted 1/4 with 0.2-μm-filtered seawater to reduce predators and increase nutrient availability for bacteria in light/dark cycles (DL), and (iv) unfiltered seawater diluted 1/4 with 30-kDa-filtered seawater to reduce predators, viruses and increase nutrient availability, in light/dark cycles (VL). The different treatments were incubated in triplicated 9L Nalgene bottles for 48h at *in situ* temperature in a water bath with circulating seawater. Light treatments were limited to photosynthetically active radiation, and dark treatments were covered with several layers of dark plastic.

Samples were taken for bacterial isolation at the same time that samples for ancillary data (reported in 37), DAPI total counts, CARD-FISH, and 16S rRNA gene amplicon sequencing. Samples were obtained at times 0 h, 12 h, 24 h, and 36 h in summer and winter or 48 h in the fall and spring experiments. For isolation, 1 mL seawater subsamples were mixed with 75 μL dimethyl sulfoxide (DMSO) in a cryovial that was stored at −80° C in triplicates. In the winter experiment, samples for isolation were not taken at 0 h, therefore 12 h after the start of the experiments was our initial time for isolation in this season.

### Community DNA extraction and sequencing

For 16S rRNA gene amplicon sequencing, samples were prefiltered through a 20 μm mesh to remove large particles and microbial biomass was concentrated onto 0.2 μm polycarbonate filters using a peristaltic pump. About 2-4 L were filtered from each replicate of all treatments. We extracted the DNA from the filters as described in Massana et al (56), and it was purified and concentrated using Amicon 100 columns (Millipore) and quantified in a NanoDrop-1000 spectrophotometer (Thermo Scientific). We stored the DNA at −80° C and an aliquot from each sample was used for sequencing using a MiSeq sequencer (2 × 250 bp, Illumina). A first run was sent to the Integrated Microbiome Resource (Halifax, NS, Canada; https://imr.bio) and a second run was sent to the Research and Testing Laboratory (Lubbock, TX, USA; http://rtlgenomics.com/) in order to improve the quality of some of the samples. Primers 515F-Y (5′-GTG YCAG CMG CCG CGG TAA) and 926R (5′-CCG YCA ATT YMT TTR AGT TT) from Parada et al (57) were used to amplify the V4-V5 regions of the 16S rRNA gene.

### Amplicon sequencing data processing and taxonomic classification

We obtained two different runs of sequences that needed to be processed separately. Initially, we used *cutadapt* (58) to trim primers. Then, ASVs were obtained running DADA2 1.18.0 version (59), which consisted in different steps. The *qscore* plots were used to inspect the quality of our samples and decide where to trim. The following step was the DADA2 process itself ran with the pool method to increase sensitivity to sequences that may be present at very low frequencies in multiple samples. Finally, we merged the two runs and removed chimeras keeping 82.7%± 0.1 sequences per sample and obtaining the final ASV table. Taxonomic assignation was performed with DECIPHER 2.16.1 version (60) at 60% confidence aligning against the SILVA database (SILVA_SSU_r138_2019.RData). Four samples with <5,000 reads were discarded, keeping a total of 308 samples for further analyses.

### Isolation of bacteria

Initial experiment sampling times (t_0_) and final (t_f_) of dark/light treatments from each season were used for isolation. Samples from 0 h of dark treatments from control and predator-reduced experiments were not employed for this purpose, since at that point there had not been time for the light regime to cause an effect, therefore the total number of samples was 42. Two different solid culture media were used, MA (Difco^TM^) and mR2A, which consisted on R2A Agar (Difco^TM^) prepared in Milli-Q water with 40 g L^−1^ Sea Salts (Sigma) (61), pH adjusted to 7.6. For liquid cultures, Marine Broth 2216 (Difco^TM^) and mR2A Broth prepared with R2A Broth (Neogen) in Milli-Q water with 40 g L^−1^ Sea Salts (Sigma) were used.

Aliquots of 100 μL of undiluted, 1:10 and 1:100 diluted seawater were spread in triplicates on agar plates and incubated at room temperature (20-25° C) until no more colonies appeared (maximum 30 days). Colonies with different morphologies were selected from each sample and streaked on new agar plates in order to obtain pure cultures. These cultures were then transferred to liquid medium, and after turbidity was detected, 100 μL from the suspensions were kept at −20° C for DNA extraction, while the rest was stored in 25% glycerol in cryovials at −80° C. Culturability was calculated as the ratio between the concentration of CFU mL^−1^ in agar plates and the total concentration of cells obtained from DAPI counts (DAPI mL^−1^, data obtained from Sánchez et al 2020).

### PCR amplification and sequencing of isolates

Genomic DNA was extracted from 100 μL of liquid cultures incubated 10 min at 99° C in a thermal cycler and 10 min at −20° C for three times. These extractions were used to PCR-amplify the nearly complete 16S rRNA gene with primers 27Fmod (5′-AGR GTT TGA TCM TGG CTC AG-3′) and 1492Rmod (5′-TAC GGY TAC CTT GTT AYG ACT T 3′) from Page et al (62). Each PCR reaction was composed of 32.75 μL Milli-Q water, 10 μL 5X Green GoTaq^R^ Reaction Buffer (Promega), 1 μL dNTPs mix (each deoxynucleotide at 10 mM), 2 μL of each primer (10μM), 0.25 μL GoTaq^R^ DNA Polymerase (Promega) and 2 μL of extracted DNA. When faint or no bands were observed on agarose gel electrophoresis following PCR, DNA extraction was repeated with DNeasy Blood&Tissue Kit (Qiagen) following the manufacturer’s recommendations. Purification and OneShot Sanger sequencing of PCR products was carried out by Genoscreen (Lille, France) with the two above-mentioned primers.

### Isolates data processing and taxonomic classification

The sequences were manually quality-checked, trimmed and assembled with Geneious software v.2022.0.1 (63). The UCLUST algorithm from USEARCH software (64) was used to cluster sequences at 99% similarity (65) in order to infer iOTUs. These iOTUs were used for all the subsequent analyses except to compare isolates and ASV sequences and to compute a phylogenetic tree to describe putative novel isolates, for which a 100% identity clustering was performed to define ziOTUs. Taxonomic classification was performed with the SINA aligner (66) against SILVA (release 138.1; 65), RDP (release 11; 66) and GTDB (release 202; 67). Additionally, isolates sequences were submitted to BLASTn v. 2.12.0+ (69) against a subset of the RDP database containing only cultured taxa (CCM) and another one containing only uncultured taxa (CEM) in order to test for their novelty. All ziOTUs that had less than 97% similarity with their CCM were considered as putatively novel strains. To assess the number of putative novel species and genera, the USEARCH software was used to cluster putative novel isolates to 97% and 94.5% similarity.

### Phylogenetic analyses

Two phylogenetic trees were constructed, one with all iOTUs to assess diversity of the complete collection, and another one only with putative novel ziOTUs to examine the phylogenetic placement of the novel strains. The general iOTUs tree was computed without reference sequences since it was only used to calculate phylogenetic diversity. For the tree with putative novel isolates, the closest sequence in SILVA Living Tree Project (LTP_12_2021; 69) for each ziOTU was inferred with BLASTn (69) and *Prochlorococcus marinus* subsp. *marinus* (ref AE017126 from LTP_12_2021) used as an outgroup. In both cases the sequences were aligned with ClustalW in Geneious software v.2022.0.1 (63) and unaligned ends were trimmed. Phylogeny was constructed using maximum-likelihood inference with RAxML-NG 0.9.0 (71) and the GTR+G+I evolutionary model. For the iOTUs tree, bootstraps converged after 1150 replications with 3% cutoff, while for the tree with putative novel isolates it did after 1450 replications with 2% cutoff. The tree with putative novel isolates was plotted with iTOL v.6 (72).

### Compositional and statistical analyses

All analyses were carried out with the R software v. R 4.1.3 (73) and RStudio software v. 1.3.1093 (74). Data manipulation was carried out mostly using packages *tidyverse* v. 1.3.1 (75) and *qdap* v. 2.4.3 (76), plots were created in *ggplot2* v. 3.3.5 (77). An iOTU table was generated with 99%-clustered isolate 16S rRNA gene sequences. Normalized and rarefied tables were used in some of the subsequent analyses. To generate these rarefied tables, an iOTU table was first grouped by treatment and then rarefied to the lowest sampling effort (80 isolates in the DL treatment at t_0_) with 1000 permutations with the package *EcolUtils* v. 0.1 (78).

We determined whether culture media, season and treatment shaped the composition of isolates at the class level. To compare iOTU composition between culture media, a proportional Venn diagram was plotted with package *VennDiagram* v. 1.7.3 (79). To estimate sampling effort in each culture media and season, rarefaction curves were performed with package *vegan* v. 2.5.7 (80) using non-normalized iOTU tables. To estimate sampling effort across treatments, rarefaction curves were inferred with both non-normalized and rarefied iOTU tables. A PCoA based on Bray-Curtis distances followed by *envfit* analysis was carried out to test β-diversity across culture media, seasons and treatments with package *vegan* v. 2.5.7. α-diversity, richness and the Chao1 estimator (81) were computed using the non-normalized iOTU table, while Shannon indices (82), FPD (83) and standarized effect size mean nearest taxon distance (84) were calculated with the normalized table. Packages *ape* v. 5.6.2 (85) and *picante* v. 1.8.2 (86) were used in order to infer FPD of the iOTUs tree. To test for differences between culture media, seasons and treatments in all these indices, ANOVA tests and Tukey’s post hoc tests were carried out with package *stats* v. 4.1.3. To test differences in culturability across culture media, Wilcoxon Rank Sum test was performed with package *stats* after rejecting normal distribution with Shapiro-Wilk Test form the same package. Welch’s T Test was used to test significant differences in proportions cultured reads between t_0_ and t_f_ of experiments (only when both samples had more than one replicate). Fisher’s Exact Test was performed to test whether some genera were more likely to be isolated in certain treatments using the non-normalized iOTU table and package *stats* v. 4.1.3. When multiple testing, p-values were adjusted with the Benjamini-Hochberg FDR method (87). All statistical tests were made with the false-discovery rate set to 0.05.

### Comparison of isolates with ASVs

The 100%-clustered full 16S rRNA gene sequences of the isolates (ziOTUs) were compared to the ASVs defined from the V4-V5 region of the 16S rRNA gene using BLASTn v. 2.12.0+ (69). Those ASVs that presented 100% similarity with one or more ziOTUs were considered as ‘cultured’. An ASV table was used to infer the rank and abundance of each cultured and uncultured taxa by treatment and season, at times t_0_ and t_f_. Additionally, the fraction of cultured taxa was calculated also for each treatment and season at times t_0_ and t_f_. The abundances used for these calculations were the means of all sample replicates. The closest neighbor to our isolates and ASVs was inferred searching our ziOTUs sequences against the NCBI 16S ribosomal RNA database (downloaded on 2022/03/01) using BLASTn. To calculate 16S amplicon sequencing abundances, corrected by their 16S rRNA gene copies, reads of each ASV were divided by their mean 16S rRNA gene copies by class except for SAR11, which was treated as a separate category. 16S rRNA gene copy numbers were obtained from rrnDB v. 5.8 (54).

### Data availability

All Supplemental Figures and Tables can be found on GitHub (https://github.com/x-rv/Manuscript-2023). The 16S rRNA gene sequences of the isolates obtained in this study were deposited in GenBank under accession numbers <u>OP342842</u> - <u>OP344484</u>. Amplicon sequencing data of the V4-V5 region of the 16S rRNA gene used in this study are publicly available in the European Nucleotide Archive under BioProject <u>PRJEB60085</u>.

## ACKNOWLEDGMENTS

This research was supported by grant RTI2018-101974-B-C22 funded by the Spanish Ministry of Economy and Competitivity and grants TM2015-70340-R and RTI2018-101025-B-100 funded by the Spanish Ministry of Science and Innovation. This work acknowledges the Severo Ochoa Centre of Excellence accreditation (CEX2019-000928-S). Xavier Rey-Velasco was supported by Spanish FI-SDUR and FPU grants.

We want to thank Núria Vigués and Verónica Melgarejo for their role in maintaining the laboratory in good shape and managing administrative procedures. We would also like to thank Mireia Burnat, Helena Catena, Adrià Auladell, Marta Sebastián, Carolina Marín-Vindas, Cèlia Marrasé, Montse Sala, Vanessa Balagué and everyone else who participated in the REMEI experiments. We also thank the Marine Bioinformatics Service (MARBITS) of the Institut de Ciències del Mar-CSIC for having provided the computing power needed to perform some of the bioinformatic analyses.

## Abbreviations

PCoA, iOTU, ziOTU, DMSO, CL, CD, PL, PD, DL, VL, BBMO, DAPI, PD, CFU, SW, RT

